# Modeling acute myeloid leukemia in a continuum of differentiation states

**DOI:** 10.1101/237438

**Authors:** H. Cho, K. Ayers, L. DePills, Y-H. Kuo, J. Park, A. Radunskaya, R. Rockne

**Affiliations:** Department of Mathematics, University of Maryland; Department of Mathematics, Pomona College; Department of Mathematics, Harvey Mudd College; Department of Hematological Malignancies Translational Science, Gehr Family Center for Leukemia Research, City of Hope; Division of Mathematical Oncology, City of Hope

**Keywords:** diffusion mapping, hematopoiesis, single cell RNA-Sequencing, developmental trajectories, nonlinear dimension reduction, cellular differentiation, acute myeloid leukemia, differentiation continuum

## Abstract

Here we present a mathematical model of movement in an abstract space representing states of cellular differentiation. We motivate this work with recent examples that demonstrate a continuum of cellular differentiation using single cell RNA sequencing data to characterize cellular states in a high-dimensional space, which is then mapped into ℝ^2^ or ℝ^3^ with dimension reduction techniques. We represent trajectories in the differentiation space as a graph, and model directed and random movement on the graph with partial differential equations. We hypothesize that flow in this space can be used to model normal differentiation processes as well as predict the evolution of abnormal differentiation processes such as those observed during pathogenesis of acute myeloid leukemia (AML).

## 1. Introduction

The recent advance of single cell RNA sequencing (scRNA-Seq) technologies has enabled a new, high-dimensional definition of cell states. There are on the order of 20,000 protein encoding genes that compose the transcriptome, which constitute a ℝ^20,000^ dimensional space. Therefore, the configuration of the transcriptome at a point in time can be represented as a coordinate vector in space. When a cell expresses genes, it “moves” in this high-dimensional gene expression phenotype space. Over time, the sequence of locations in the space of a given cell creates a trajectory. Dimension reduction techniques are commonly used to map the larger space into a lower dimensional space, for instance, ℝ^2^ or ℝ^3^, at which point the cells are clustered based on a similarity metric and recategorized. This process has revealed a continuum of cell phenotypes, with intermediate states connecting canonical cell states. The most prominent example of this process is in hematopoietic cell differentiation.

Normal hematopoiesis is long thought to occur through stepwise differentiation of hematopoietic stem cells following a tree-like hierarchy of discrete multipotent, oligopotent and then unipotent lineage-restricted progenitors (Figure 1A). The classical model of hematopoiesis considers differentiation as a stepwise process of binary branching decisions, famously represented as a potential landscape by Waddington (Waddington 1957). However, this model is based on bulk characterization of prospectively purified immunophenotypic cell populations. Recent advances in scRNA-Seq technologies now allow resolution of single cell heterogeneity and reconstruction of differentiation trajectories which have been applied to a number of different cellular systems, from hematopoiesis to breast endothelial cell differentiation (Hamey et al. 2016; Velten et al. 2017; Bach et al. 2017; Nestorowa et al. 2016). These efforts have led to the new view that hematopoietic lineage differentiation occurs as a continuous process, which can be mapped into a continuum of cellular and molecular phenotypes (Figure 1B). Hematopoietic malignancies arise from dysregulated differentiation and proliferation of hematopoietic stem cells and progenitor cells upon accumulation of oncogenic genetic mutations and/or epigenetic alterations. Therefore, characterizing disordered hematopoiesis based on discretely defined phenotypic populations has been sometimes challenging. It is now possible to view pathologic hematopoiesis through a continuum of cellular and molecular phenotypes and capture the heterogeneity, differentiation plasticity and dysregulated gene expression associated with malignant transformation.

**Figure 1.**
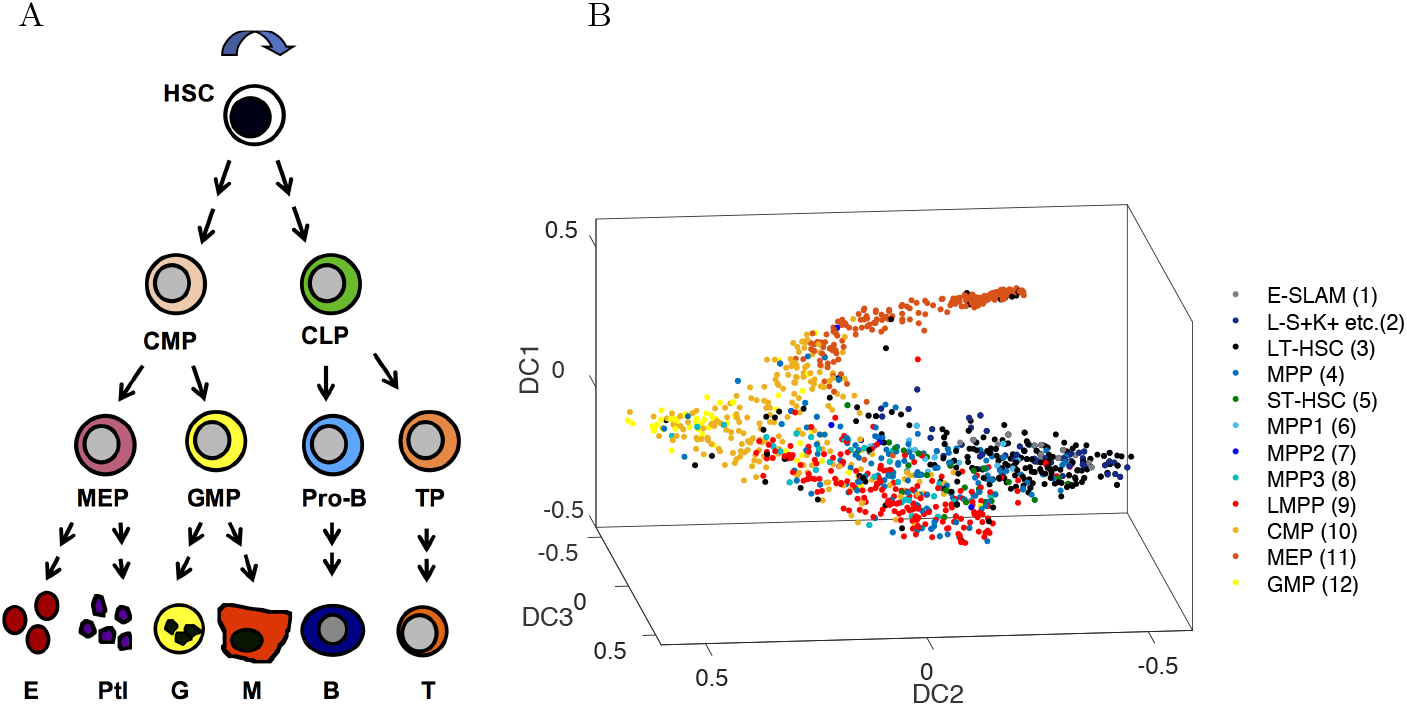
A) Classic representation of a linear hierarchy of discrete cell states, from hematopoietic stem cell (HSC) to committed common myeloid progenitor (CMP) and common lymphoid progenitor (CLP) cells, on down to terminally differentiated cells such as platelets, macrophages, B and T cells. B) The classical view is contrasted with a nonlinear continuum representation of hematopoietic cell differentiation states using diffusion map dimension reduction of scRNA-Seq data (figure recreated from data available in Nestorowa et al. (2016)). Colors representing cell identities in A) and B) are coordinated. Cell types in B) are a subset of cells represented in A).

This new view of biology forces us to reconsider the mathematical approaches we use to model cell states and behaviors. Instead of building mathematical models which identify discrete cell populations and assign mathematical rules for their evolution and interactions, we may now consider a continuum of cellular states, and model movement between these states in aggregate as a flow of mass on a structured graph. Modeling differentiation in this manner reduces the number of parameters and thus the complexity of the mathematical model by representing many cell populations and states in a single variable. At the same time this increases biological resolution of the system by characterizing an infinite number of sub-states in a continuum representation. Here we consider a model of hematopoietic cell differentiation and associated disorders as a flow and transport process in a continuous differentiation space as a test system for a more general approach of modeling the temporal evolution of a continuum of cell states.

This manuscript is structured as follows: first, we review the state of the art of dimension reduction methods that are used to construct and define hematopoietic differentiation spaces that can be represented as graphs, including a review of Schien-binger et al.’s method for modeling transport on a graph from reduced dimension gene expression data (Schiebinger et al. 2017). Then we introduce our partial differential equation (PDE) model of flow and transport on a graph, and illustrate the model on simple “Y” shaped graph. We then calibrate our model to a graph constructed from publicly available scRNA-Seq data of normal hematopoiesis. Finally, we use our model to simulate abnormal hematopoietic cell differentiation processes observed during the evolution of acute myeloid leukemia (AML), a form of aggressive hematologic malignancy. We conclude with a brief discussion of the limitations and potential future applications of this modeling approach.

## 2. Construction of a differentiation continuum

While the focus of this paper is not dimension reduction techniques or *pseudotime* reconstruction, we begin by summarizing a some of these techniques that are most relevant to our modeling approach, without advocating for one over another. The relationship between time and pseudotime within a mathematical model of cell differentiation is analogous to the relationship between age structured and stage structured models in ecology. Cell differentiation data yield information about cells at various stages of differentiation, but generally do not provide time-specific data. A pseudotime model is one that considers the differentiation stage of a cell population instead of the time in which a cell is in a certain state.

A broad range of techniques have been developed to provide insight into interpretation of high dimensional data. These techniques provide a first step in our approach to modeling the evolution of cell states in a continuum and play a critical role in characterizing differentiation dynamics. Each method has its own advantages and disadvantages. As a result, application of different data reduction, clustering methods, and pseudotime ordering on the same data set will produce different differentiation spaces on which to build a dynamic model. We will use one particular dimension reduction approach as an example, but our framework allows one to select from a variety of approaches. In this section we will provide a brief description of a subset of such techniques.

### 2.1. Dimension reduction techniques

Several techniques have been developed to interpret the high-dimensional differentiation space, including principal component analysis (PCA), diffusion mapping (DM) and t-distributed stochastic neighbor embedding (t-SNE). Each of these methods project data into a lower dimensional space. As discussed in this section, different techniques provide different shapes and differentiation spaces, and so some techniques are better suited to certain data sets than others. For instance, one commonly used dimension reduction technique is principal component analysis (PCA), a linear technique. While PCA is computationally simple to implement, the limitation of this approach lies in its linearity - the data will always be projected onto a linear subspace. If the data shows a trend that do not lie in a linear subspace—for instance, if the data lies on an embedding of a manifold in Euclidean space—this trend will not be captured with PCA.

In contrast, diffusion mapping (DM) and t-stochastic neighbor embedding (t-SNE) are two non-linear dimension reduction techniques. Diffusion maps apply a diffusion operator to the data set, and determine embeddings of the data in a lower dimensional Euclidean space by considering eigenvalues and eigenvectors of this operator. In Haghverdi, Buettner, and Theis (2015), the authors propose and adapt diffusion maps for single-cell data sets, and further claim that diffusion maps are better suited for RNA-Seq data than either PCA or even other non-linear techniques, including t-SNE. t-SNE, meanwhile, is a machine learning dimension reduction technique that is particularly good at mapping high dimensional data into a two or three dimensional space, allowing for the data to be visualized in a scatter plot. t-SNE aims to find a map from the data set to two or three dimensional Euclidean space that minimizes the Kullback-Leibler divergence between the two probability distributions in the original and reduced space. This optimization problem is often solved using gradient descent methods.

In van Unen et al. (2017), a new technique for examining high dimensional mass cytometry data, known as hierarchical stochastic neighbor embedding (HSNE) is presented. Mass cytometry allows for the examination of several cellular markers on samples made up of vast quantities of cells. These data sets are difficult to examine statistically due to their large size and high dimensionality. Therefore, due to all these factors, pre-existing dimension reduction techniques are not optimal for mass cytometry data. What makes HSNE useful is that it preserves local data structure while allowing examination of the full data set. HSNE ultimately constructs a hierarchy of subsets of the dataset 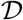:

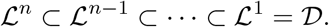

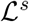 is the set of landmarks representing the data at scale *s*. The algorithm begins by considering a Markov Chain on 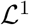 from a k-nearest neighbor graph. We then consider the invariant distribution of this Markov Chain, and points in 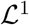 that have a probability in the invariant distribution above some cutoff value are selected to be in 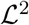. For each set of landmarks 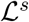, for *s* > 1, an “area of influence” (AoI) is determined as well. This is done by considering a set of random walks started from each landmark in 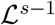. Once a random walk reaches a landmark in 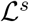, it terminates. The AoI for each landmark is the set of all landmarks in 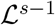 that reached this landmark at least once. The extent to which the AoIs of two landmarks in 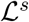 overlap determines their similarity, and this defines a new neighborhood graph on 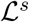. The process is then repeated by considering a Markov chain on this graph. This construction allows for the creation of a hierarchy of different neighborhoods as given by the resulting similarities. These neighborhoods are then examined in a backwards order, from coarsest to finest, and can be embedded in a lower dimensional space using Barnes Hut Stochastic Neighbor Embedding (BH-SNE), a variant of tSNE. Starting with a certain subset 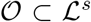 of landmarks, the user is able to “drill in” the data by selecting a subset 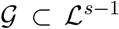. Landmarks thus can be added on demand according to the level of detail required by the user - thus, HSNE is an approach that is useful for data that requires different levels of detail at different scales.

Each of these dimension reduction methods has strengths and weaknesses depending on the question(s) being asked of the data. Moreover, each method will produce a distinctly different shape in the lower dimensional projection. Therefore, the choice of dimension reduction technique is a critical step in analyzing any high-dimensional data set. For the purpose of analyzing cell transition probabilities and inferring trajectories within the reduced space, Nestorowa et al. (2016) and others have chosen to use diffusion mapping to analyze cell differentiation.

### 2.2. Pseudotime ordering of differentiation states

For data without temporal information, pseudotime methods are available to infer a sequence of biological states from single time point data. Diffusion mapping can be used to infer a “diffusion pseudotime” (Haghverdi et al. 2016; Nestorowa et al. 2016). In particular, Haghverdi et al. (2016) develops an efficient diffusion pseudotime approach by rescaling the diffusion components by a weighted distance in terms of the eigenvalues, derived by considering a random walk according to a transition matrix that specifies the probability of transitioning from any single cell to another in an infinitesimal amount of time. Alternative pseudotime approaches include Wishbone (Setty et al. 2016) that uses shortest paths in a k-nearest neighbor (kNN) graph constructed in diffusion component space to construct an initial ordering of cells, TASIC (Rashid, Kotton, and Bar-Joseph 2017) that is able to incorporate time information and identify branches and incorporate time information in single cell expression data by considering it as developmental processes emitting expression profiles from a finite number of states, and Monocle (Qiu et al. 2017b,a) that fits a principal graph (Mao et al. 2015) and uses a reversed graph embedding technique which simultaneously learns a low dimensional embedding of the data and a graphical structure spanning the dataset.

Finally, when the data are collected at multiple time points, the transition rates between the nodes can be obtained after partitioning the cell data. For instance, Schiebinger et al. (2017) employs graph clustering (Levine et al. 2015; Shekhar et al. 2016) and optimal transport methods to understand the dynamics in the reduced space of cell data. We describe the optimal transport (OT) method in an effort to provide a clear distinction between the OT algorithm and our modeling approach.

### 2.3. Optimal transport

Schiebinger et al. (2017) have proposed a model and algorithm for constructing a graph in differentiation space, oriented in pseudotime given temporal data. This method is not a dimension reduction technique, rather, the optimal transport algorithm is applied to a time series of reduced dimension single cell gene expression profiles. The time series is made up of a sequence of samples at different times *t_i_* for *i* ∈ {1, …, *n*} from some distribution on ℝ^*n*^. An distribution 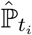 is defined by each sample *S_i_*:

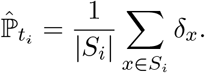

Together, as a sequence, these inferred distributions 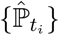 form what is known as an empirical developmental process. The goal is then to determine, as closely as possible, what the true underlying Markov developmental process ℙ_*t*_ is by examining what are known as transport maps between pairs 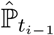 and 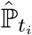. A transport map for a pair 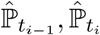 is a distribution *π* defined on ℝ^*n*^ × ℝ^*n*^ such that 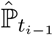 and 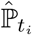 are the two marginal distributions of *π*. Thus, given a function *c*(*x, y*) that represents the cost to transport some unit mass from *x* to *y*, the goal is to minimize

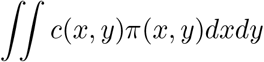

subject to

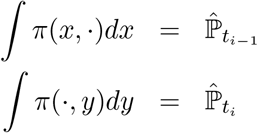

Schienbinger et al. further refine this algorithm by including a growth term in their transport plan to allow for cellular proliferation between time points. The classical optimal transport algorithm is formulated with conservation of mass in mind. Optimal transport can thus be used to estimate the ancestors and descendants of a set of cells. Cells are clustered using the Louvain-Jaccard community detection algorithm on the reduced dimension gene expression data in 20 dimensional space. Schienbinger et al. thus identified 33 cell nodes, which were then used as starting populations from which developmental trajectories could be analyzed. These can be thought of as nodes on a graph visualized with force-directed layout embedding, and edges represent the motion in pseudotime.

## 3. Modeling on the differentiation continuum

To illustrate our modeling technique, we assume that we have constructed a simple branched manifold or graph situated in the differentiation space. This graph is not a set of discrete nodes, rather, the graph and its edges represent a continuum of canonical states and intermediate states of differentiation. Assuming that the graph and the temporal evolution on the graph is obtained by any one of the various data analysis techniques summarized in Section 2 including optimal transport (Schiebinger et al. 2017), diffusion pseudotime methods (Haghverdi et al. 2016; Nestorowa et al. 2016; Haghverdi et al. 2016), Wishbone (Setty et al. 2016), TASIC (Rashid, Kotton, and Bar-Joseph 2017), and Monocle (Qiu et al. 2017b,a), we develop a PDE model that describes the dynamics in this differentiation continuum.

### 3.1. PDE model on a graph

Let us define the graph *G* obtained in the reduced space. The node set of *G* is denoted as 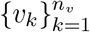, where *n_v_* is the total number of nodes, and the edge of the graph connecting in the direction from the *i*-th to the *j*-th node as *e_ij_*. We also introduce an alternate description of the graph with respect to the edge, that is more convenient for describing the PDE model. If the total number of nontrivial edges is *n_e_*, we take 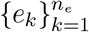 with the index mapping 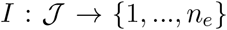 on the set of nontrivial edges 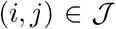, and the end points in the direction of cell transition as 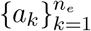 and 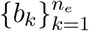, respectively. We remark that 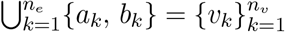.

Then, the cell distribution *u*(*x, t*) on graph *G* can be modeled by an advection-diffusion-reaction equation (Evans 2010) on the edges as

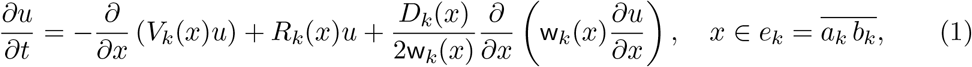

where *x* is a one dimensional variable parameterized on each edge *e_k_* from *a_k_* to *b_k_*. The advection coefficient *V_k_*(*x*) models the transition between the nodes and the obtained transition rate per unit time can be interpolated between the nodes, that is,

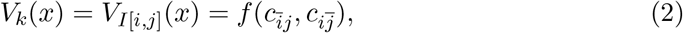

where *f* is an interpolation function, and 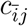 and 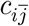 the flow rate near the node *i* and *j*.

Cell proliferation and apoptosis can be modeled by the reaction coefficient *R_k_*(*x*). In addition to natural proliferation and apoptosis, this term can also model abnormal tumorous cell growth or the effect of targeted therapy by localized Gaussian or Dirac-delta functions centered at the location of the cell type on the graph.

The diffusion term represents the noise or instability of the cell landscape. It involves two parameters *D_k_*(*x*) and w_*k*_(*x*) describing the noise in the direction parallel and vertical to the edge, respectively. Considering the cell density as a random process subject to isotropic white noise with magnitude σ, the diffusion term becomes *D_k_* = *σ*^2^ and w**k** = 1 (Evans 2014). On the other hand, if the noise is anisotropic, we employ the procedure of obtaining the PDE model on a graph as a projection from PDE defined on narrow domains (Cerrai and Freidlin 2017; Freidlin and Hu 2013). This is valid when the cell data in the reduced space lies in one dimensional narrow channels. While *D_k_*(*x*) represents the diffusion along the edge, w_*k*_(*x*) is the width of the narrow channel in the perpendicular direction to the projection. Finally, we remark that all the coefficients can be time dependent.

In addition to the governing equation on the edges, the boundary condition at the nodes are critical when describing the dynamics on the graph. The boundary condition on the cell fate PDE model can be classified into three types, the initial nodes that do not have inflow 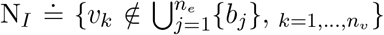, the final nodes without outflow 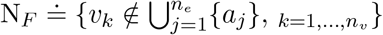, and the intermediate nodes,

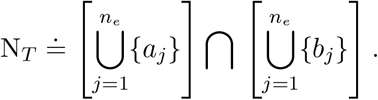

On the intermediate nodes *v_k_* ∈ N_*T*_, mixed boundary conditions can be imposed to balance the cell inflow and outflow as

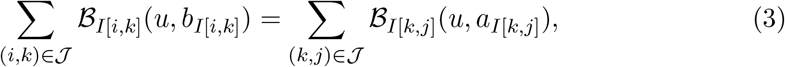

where 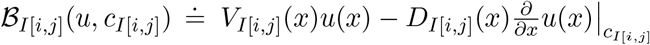 with continuity conditions as Dirichlet boundary conditions

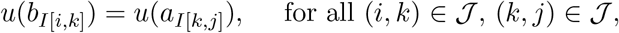

for a fixed *k*. The cell outflow boundary conditions on the final nodes *v_k_* ∈ N_*F*_ are imposed as reflecting boundary conditions

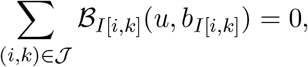

and *u*(*b*_*I*[*i, k*]_) = *u*(*b*_*I*(*j,k*]_) for all (*i, k*) and (*j, k*) in 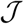. Similarly this can be imposed on the initial nodes *v_k_* ∈ N_*I*_ as 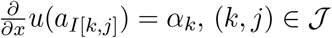 or *u*(*a*_*I*[*k,j*]_) = *α_k_*, 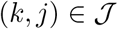 depending on whether the model describes the cell inflow flux or a proscribed inflow density.

#### 3.1.1. Example on a Y shaped graph

To illustrate our approach, we derive the PDE model given in Equation (1) on a simple Y shaped graph. This example is motivated by supposing that the cells have two different cell fates with one branching. For instance, assume that the cell data projected onto the first two diffusion components, `DC1` and `DC2`, are as in Figure 3A and the temporal direction is from left to right, as indicated by the arrows in the figure. We define the Y shaped graph with four nodes *v*_1_ = (−1, 0), *v*_2_ = (0, 0), *v*_3_ = (1, 1), and *v*_4_ = (1, −1), and three edges 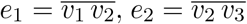, and 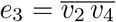. For simplicity, let us assume that the edges are straight lines and parametrize the variables on each edge as *e*_1_(*x*) = (*x* − 1, 0), *e*_2_(*x*) = (*x, x*), and *e*_3_(*x*) = (*x, −x*), so that *x* ∈ [0, 1]. When there is possibility for confusion, we use subscripts on the *x*-variables to specify which edge is parametrized. So, for example, *x*_2_ parametrizes *e*_2_. Then, the PDE model on each parametrized edge can be written as

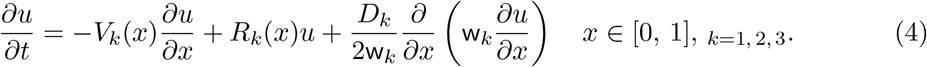

**Figure 2.**
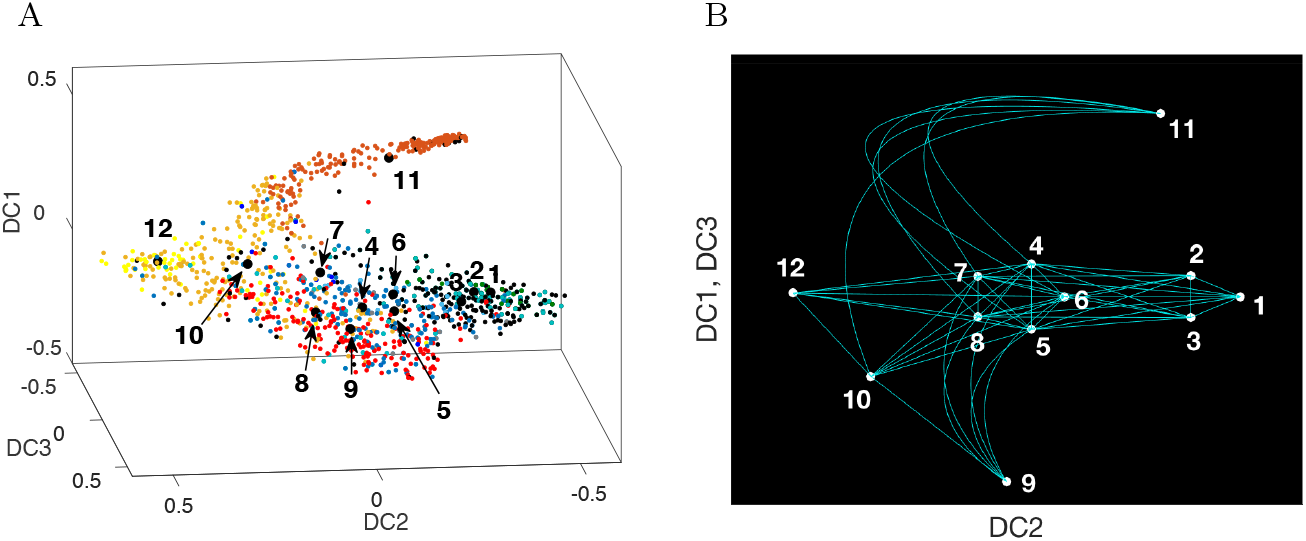
A) For the 12 cell types identified in Nestorowa et al. (2016), the center of mass of each cell type is used to define a node on an abstracted graph B). Edges between nodes are constructed based on inferred trajectories on the graph based on diffusion pseudotimes starting from nodes 1, 2, and 3, to nodes 4-8, then to the progenitor nodes 9-12. The graph represents a continuum of cell states (edges) that includes identification of canonical cell states along the continuum (nodes 1-12) (Table 1).

**Figure 3.**
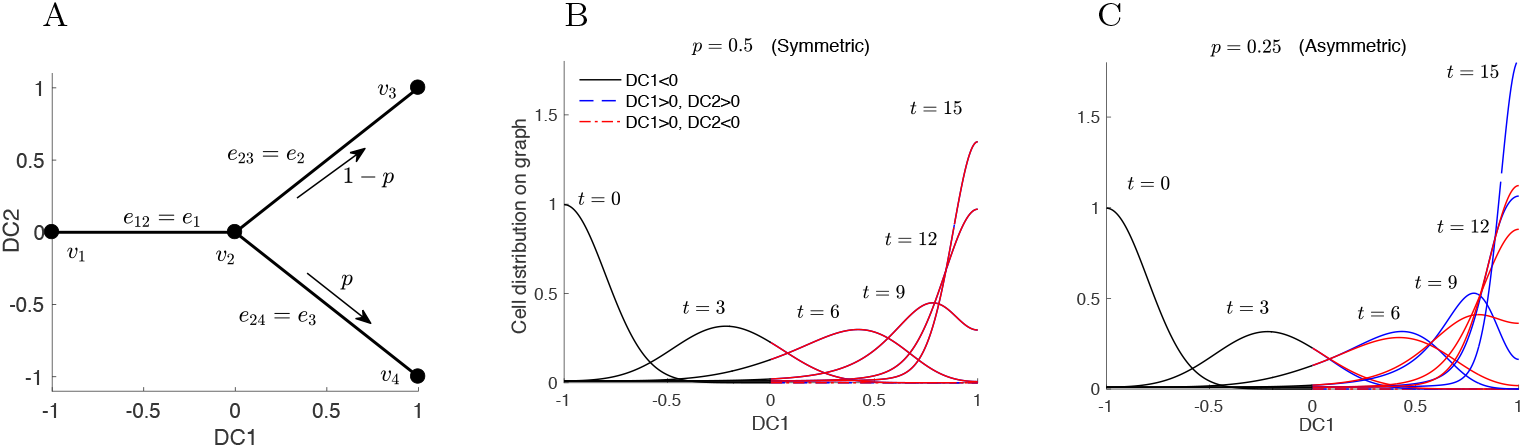
We use a simple “Y” shaped graph to illustrate our model. A) The graph is defined by four nodes 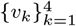 and three edges *e*_1_ = *e*_12_, *e*_2_ = *e*_23_, and *e*_3_ = *e*_24_ within two components of a diffusion map (`DC1, DC2`). The transfer rate from *v*_2_ to *v*_3_ and *v*_4_ is taken as 1 − *p* and *p*, respectively. B) The evolution of the cell density solution from the initial condition (*t* = 0) concentrated at the left end, `DC1` = −1, to a density concentrated at the right ends, `DC1` = 1, at *t* = 15. In the symmetric case, *p* = 0.5, the two branches evolve in the same way; C) in the asymmetric case, *p* = 0.25, the cell density is larger at *t* =15 on the upper branch, shown in blue, compared to the lower branch, shown in red.

We consider the case that the cells transfer from *v*_1_ to *v*_2_ in *n_T_* unit time, differentiate into each cell type with proportion *p* and 1 − *p*, and accumulate at `DC1` = 1, where the cells are fully differentiated. Then,

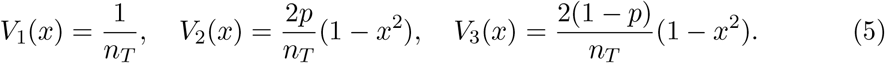

Also, we assume that the differentiation process is subject to random white noise with a constant magnitude *σ* so that *D_k_* = *σ*^2^, and the width of the cell located area in the vertical direction is constant regarding the differentiated ratio, that is, w_1_/2 = w_2_ = w_3_ is a constant. Figure 3 plots the two examples of symmetric differentiation *p* = 0.5 and asymmetric differentiation *p* = 0.25.

Finally, proliferation of the stem cells with cell cycle *λ* and apoptosis of the differentiated cells with rate *d_k_* can be modeled as

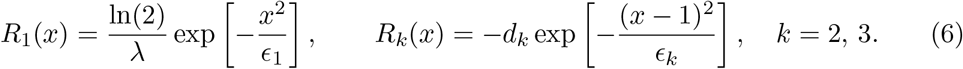

The boundary type of the nodes are classified, according to our description above, as N_*I*_ = {*v*_1_}, N_*T*_ = {*v*_2_}, and N_*F*_ = {*v*_3_, *v*_4_}. Thus, we impose the *gluing* boundary condition at *v*_2_, using subscripts on the parametrization variables to denote the specific edge, as

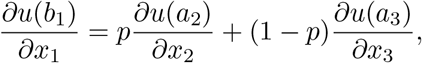

with *u*(*b*_1_) = *u*(*a*_2_) = *u*(*a*_3_), a constant inflow boundary condition at *v*_1_, and reflecting boundary conditions at the end nodes *v*_3_ and *v*_4_,

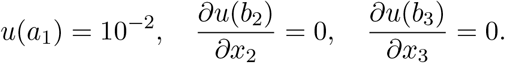

Simulations of this simple model are shown in Figure 3, where densities on edges *e*_2_ and *e*_3_ are plotted in different colors. We see that an initial density concentrated near *v*_1_ moves to the right, branches at *v*_2_, and becomes absorbed at *v*_3_ and *v*_4_. In the symmetric case, when *p* = 0.5, the density is the same on each of the two branches to the right of *v*_2_, so that the two curves are plotted on top of each other. When *p* = 0.25, the density profile is not symmetric: more cells move along the upper branch than on the lower branch.

## 4. Simulation results

In this section, we employ the framework developed in section 3.1 to the mouse hematopoietic stem and progenitor cell data in Nestorowa et al. (2016).

### 4.1. Model of normal adult hematopoiesis

To calibrate our model, we first apply it to normal hematopoietic cell differentiation trajectories identified in Nestorowa et al. (2016). Nestorowa et al. characterize early stages in hematopoiesis with twelve cell types, shown in Table 1 and Figure 2, including E-SLAM (CD48-CD150+CD45+EPCR+), long-term HSCs (LT-HSCs), shortterm HSCs (ST-HSCs), lymphoid-primed multipotent progenitors (LMPPs), multipotent progenitors (MPPs), and megakaryocyte-erythroid progenitors (MEPs), common myeloid progenitors (CMPs), and granulocyte-macrophage progenitors (GMPs). We consider these twelve cell types as the twelve nodes, *v_k_*, in our graph, and add 51 edges to model the hematopoietic cell hierarchy (see Figure 1A) and pseudotime computed in Nestorowa et al. (2016) (see Figure 4A). This graph represents a continuum of canonical and intermediate states of hematopoietic differentiation with nodes and edges, respectively. The spatial variable in our PDE model represents the differentiation state of the cell.

**Figure 4.**
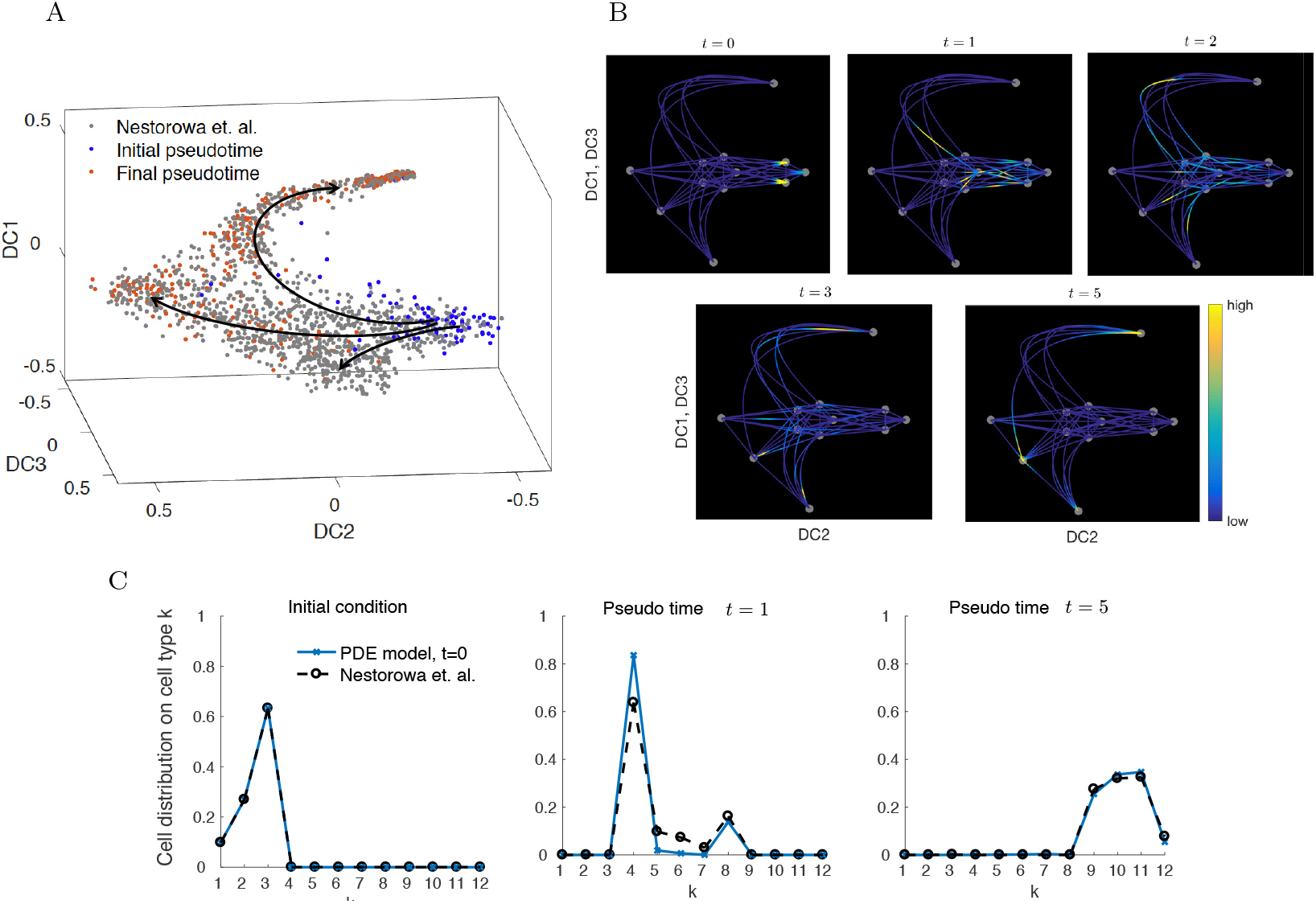
A) The cell data colored by pseudotime analysis produced by the Wanderlust algorithm applied to data mapped to diffusion space in Nestorowa et al. (2016). The initial point in pseudotime is taken from the HSC cells and the final pseudotime from the progenitor cells. B) Cell distribution computed by the PDE model describing normal conditions on the graph from *t* = 0 to *t* = 5. The cells flow from E-SLAM and LT-HSC nodes on the right to the LMPP, CMP, MEP, and GMP nodes on the other three ends (bottom, top, and left), following the pseudotime trajectories identified in A). C) Comparison of the cell type distribution computed by the PDE model described in Equation (1) and the reference data from Nestorowa et al. (2016). The reference distribution (Nestorowa et al.) is computed by clustering the initial, middle and final pseudotime cells from A) into 12 cell nodes. By considering *t* = 5 as the final pseudotime in the PDE model, the values of the solution at the nodes agree well with the reference data.

**Table 1.** Index of cell identities and labels. Long- and short-term hematopoietic stem cells (LT-HSC, ST-HSC); multipoent potent progenitors (MPP), lymphoid-primed multipotent progenitors; common myeloid progenitors (CMP); megakaryocyte-erythroid progenitors (MEP); granulocyte-macrophage progenitors (GMP).

| Cell identities and labels |  |  |  |
| --- | --- | --- | --- |
| ID | Cell type | ID | Cell type |
| 1 | E-SLAM | 7 | MPP2 |
| 2 | L-S+K+ CD34-<br>Flk2+ CD48- CD150+ | 8 | MPP3 |
| 3 | LT-HSC | 9 | LMPP |
| 4 | MPP | 10 | CMP |
| 5 | ST-HSC | 11 | MEP |
| 6 | MPP1 | 12 | GMP |

The colored and labeled clustered cell data and the corresponding graph are shown in Figure 2. The location of the nodes on the graph is not chosen to be identical to the data, but for an illustrative purpose to represent `DC2` and `DC1`/`DC3`. The edges are chosen according to the pseudotime progression from the E-SLAM and HSCs (nodes 1-3) to the progenitor cells (nodes 9-12).

The parameters of the PDE model of cell differentiation under normal conditions are chosen to reproduce the distribution of cell types from Nestorowa et al. (2016) at the initial and final pseudotime (Figure 4C). The reference distribution is computed by counting the relative number of cells in each cluster at the initial and final pseudotime. The initial and final cell distribution is concentrated on nodes 1-3 and 9-12, respectively. The distribution of cells in the remaining states, represented by nodes 4-8, goes from 0 at time *t* = 0 to positive at time *t* = 1, and back to 0 at *t* = 5, the end of the simulation. We remark that the ratios of the number of cells in each node are used to compute the advection coefficients *V_k_* in Equation (2), where we take the drift *V*_*I*[*i,j*]_ from cell type *i* to another other cell types *j* to be proportional to the ratio plotted in the figure. For instance, the outflow from *v*_5_ to nodes 9-12 is taken to be proportional to the reference distribution at the final pseudo time in Figure 4C. The diffusion coefficient is taken as *D_k_* = 10^−2^ within the subsets of nodes {1, 2, 3}, {4, …, 8}, {9, …, 12}, and *D_k_* = 10^−3^ between these subsets of nodes. This selection of diffusion coefficients takes into consideration the distance between the center of mass of the node subsets in the diffusion space. We assume a constant proliferation rate of *R_normai_* ≐ 0.956 for all edges.

For the implementation, we discretize the system using a fourth-order finite difference method and 100 grid points on each edge, and a third-order Runge-Kutta method in time with time step 10^−4^. Figure 4C compares the solution to the PDE in the normal condition to the reference distribution. The initial condition of the PDE is taken as the initial reference distribution, and we compute the solution up to time *t* = 5. The solution at *t* = 5 is similar to the reference distribution at final pseudotime. Also, the solution at *t* = 1 is close to the distribution of the remaining cells excluding the initial and final cells. Figure 4B shows the cell distribution on the graph from time *t* = 0 to *t* = 5. We observe that the support of the cell density shifts from the initial nodes 1-3, representing HSCs, to nodes 9-12, representing progenitor cells.

### 4.2. Acute myeloid leukemia (AML)

With our model calibrated to data obtained from normal hematopoietic differentiation trajectories, we now simulate AML pathogenesis. AML results from aberrant differentiation and proliferation of transformed leukemic stem cells (LSC) and abnormal progenitor cells. Parallel to the hierarchy of normal hematopoiesis, malignant hematopoiesis has also been considered to follow a hierarchy of cells at various differentiation states although with certain levels of plasticity (see Figure 5).

**Figure 5.**
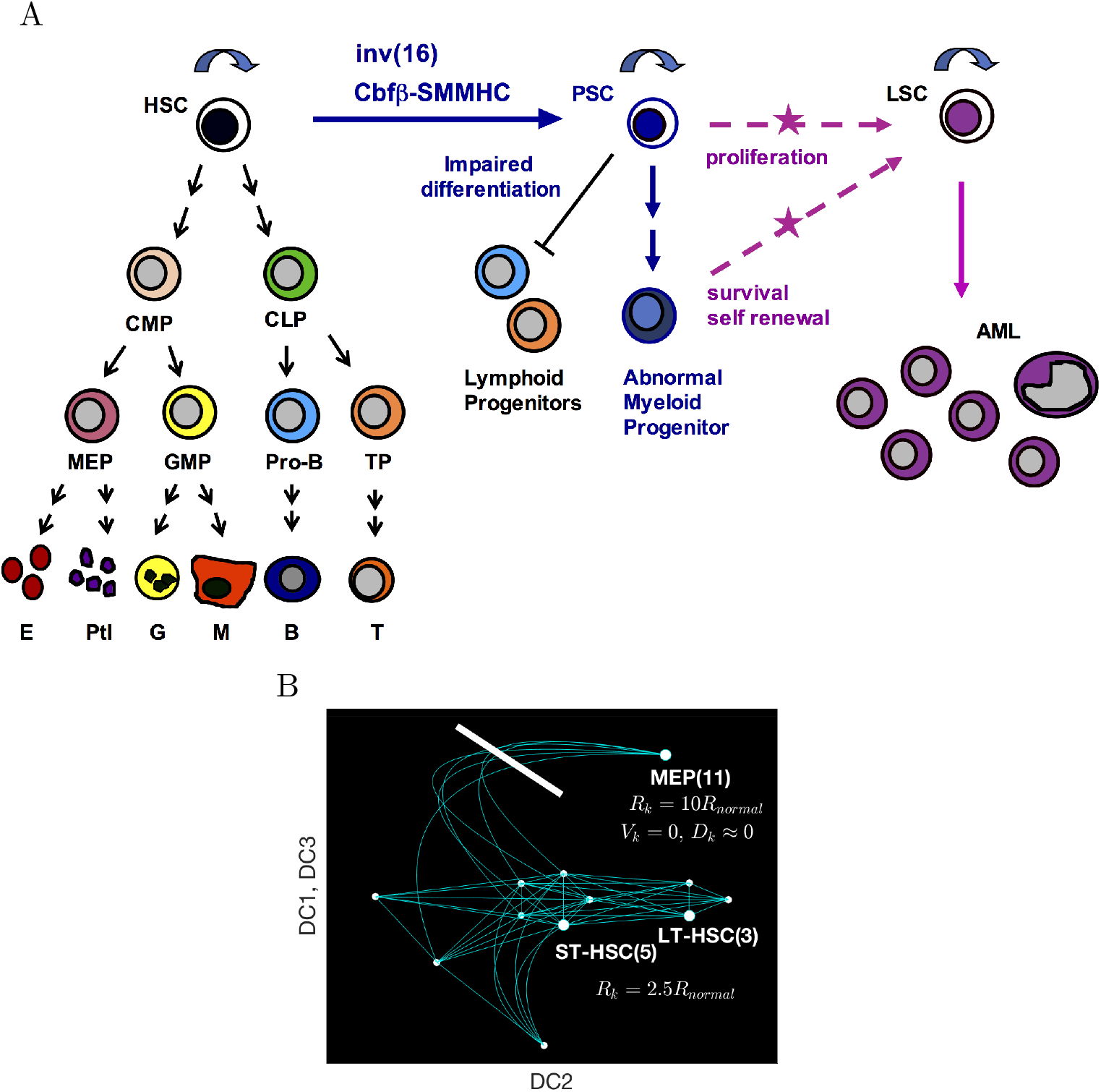
A) Acute myeloid leukemia (AML) is a cancer of aberrant differentiation and proliferation of partially transformed preleukemic stem cells (PSC), fully transformed leukemic stem cells (LSC) and abnormal progenitor cells. Malignant hematopoiesis is also considered to follow a hierarchy of cells at various differentiation states, although with higher levels of plasticity. B) Schematic illustration of AML pathogenesis in the differentiation continuum. To simulate inv(16) driven AML, the proliferation *R_k_* connecting the nodes 3, 5, and 11 is increased and the flow toward the node 11, *V_k_* and *D_k_*, is blocked.

Given the aberrant differentiation and plasticity associated with the pathology of AML, modeling in a continuous differentiation space offers the advantage over discrete models that all pathological and intermediate cell states can be captured. Here, we model the progression of AML using a genetic knock-in mouse model that recapitulates somatic acquisition of a chromosomal rearrangement, inv(16)(p13q22)(Liu et al. 1993, 1996), commonly found in approximately 12 percent of AML cases. Inv(16) rearrangement results in expression of a leukemogenic fusion protein CBF *β*-SMMHC, which impairs differentiation of multiple hematopoietic lineages at various stages (Castilla et al. 1999; Kuo et al. 2006; Kuo, Gerstein, and Castilla 2008). Based on previous studies using the inv(16) AML mouse model (Kuo et al. 2006; Cai et al. 2016), we know that expression of CBFβ-SMMHC leukemogenic fusion protein induces moderate expansion of preleukemic stem cells and abnormal myeloid progenitors skewed towards a MEP-like or pre-megakaryocyte/erythroid (Pre-Meg/E) phenotype (Figure 5A).

To simulate AML pathogenesis, we increase the proliferation rate of MEP (node 11) by 10 times, that is, *R*_*I*[*i*,11]_ = 10*R_normal_*, while blocking the flow to the MEP by taking zero advection coefficient on the edge that is connected to *v*_11_, i.e., *V*_*I*(*i*,11)_ = 0. We also lower diffusion to *D*_*I*(*i*,11)_ = 10^−5^ and linearly increase it to the original value to model the imperfect differentiation block involved in AML pathogenesis. In addition, the proliferation rate of LT-HSC and ST-HSC (nodes 3 and 5) is increased by 2.5 times by *R*_*I*[3,*j*]_ = *R*_*I*[*i*, 3]_ = *R*_*I*[*i*,11]_ = *R*_*I*[11,*j*]_ = 2.5*R_normal_*. This is illustrated in Figure 5B.

Figure 6 shows the total number of cells in each cell type in normal condition and AML condition starting at *t* = 3. In normal condition, the CMP, MEP, and LMPP cells dominates the population after *t* ≥ 3. However, in the AML case, the MEP cells increases up to 100 times of the normal condition and dominates the population. Figure 6C plots the number of cells in each cell type separately, where we can observe the increasing number of cells not only in MEP, but also the intermediate cell types, 4-8. Figure 7 compares the cell distribution on the graph between the normal and AML case at time *t* = 8. In the AML case, the peak is shown on the edges near MEP cells. The continuum of intermediate cell types, represented as numbers of cells along the edges of the graph are plotted in Figure 8. The cell distribution in the normal case at *t* = 1 and *t* = 3 shows the cell population moving on the edges from HSCs to progenitors states. Under normal hematopoeisis, we observe the flow of cells along the continuum from a stem like state to a progenitor state, with an even distribution of all types of progenitor cells. However in the AML case, we predict the emergence of novel intermediate cell types, including a mixed CMP-MPP3 and CMP-MEP cell type, in addition to a 10 fold increase in the number of canonical MEP cells. These indeterminate cells may exhibit phenotypic properties of both cell types on either side of the edge (node *i* and/or node *j*). This cell state may be unstable, phenotypically plastic, may be in an abnormal state or process of differentiation, or perhaps even undergoing a selection pressure to induce transformation.

**Figure 6.**
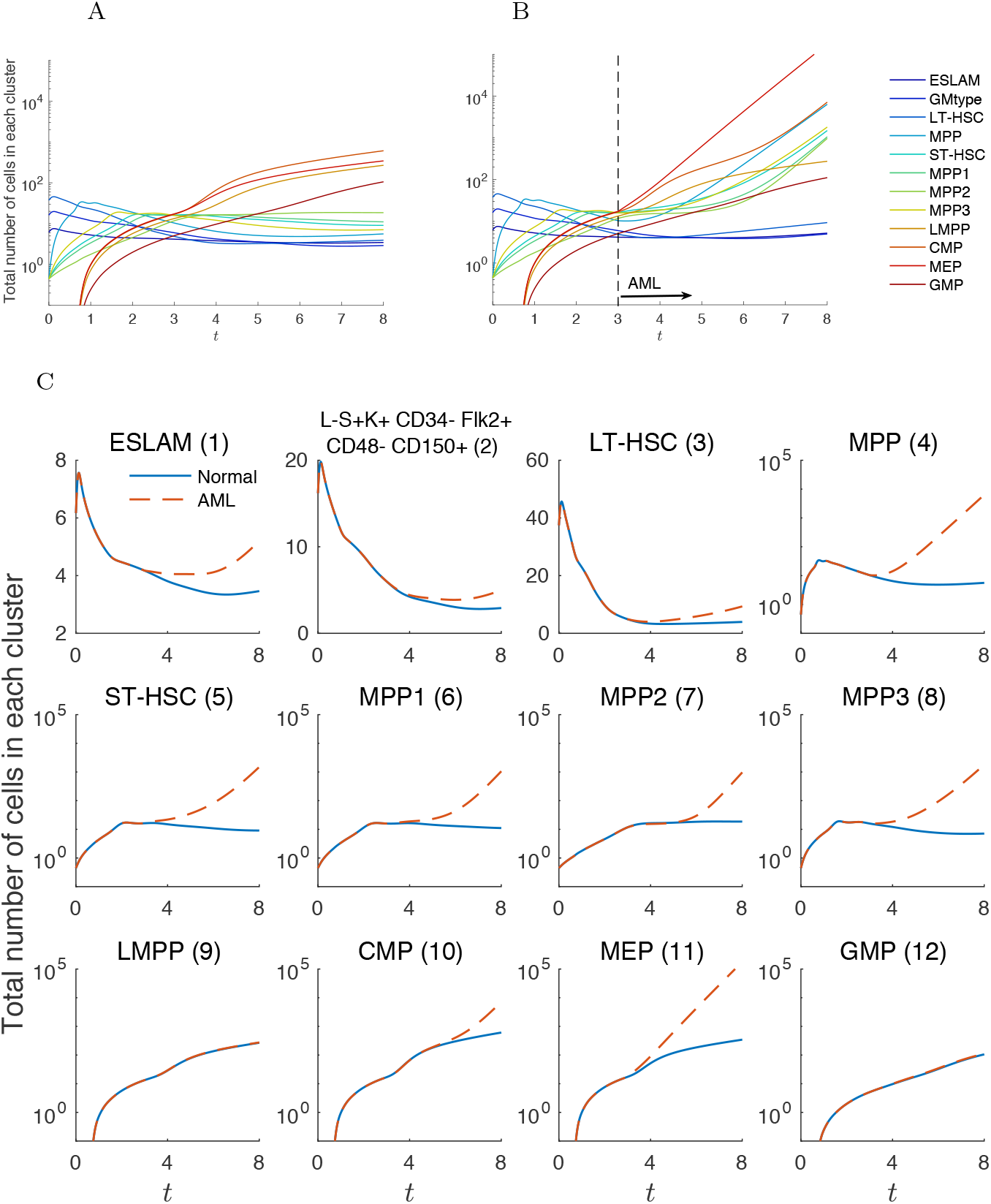
Total number of cells in each node up to *t* = 8 in A) normal condition and B) AML pathogenesis. The AML simulation is started at *t* = 3. Compared to the normal case, cells in MEP, LT-HSC, and ST-HSC increase as well as other cell types. Figure C) compares the number of cells between the normal and AML case for each cell type individually.

**Figure 7.**
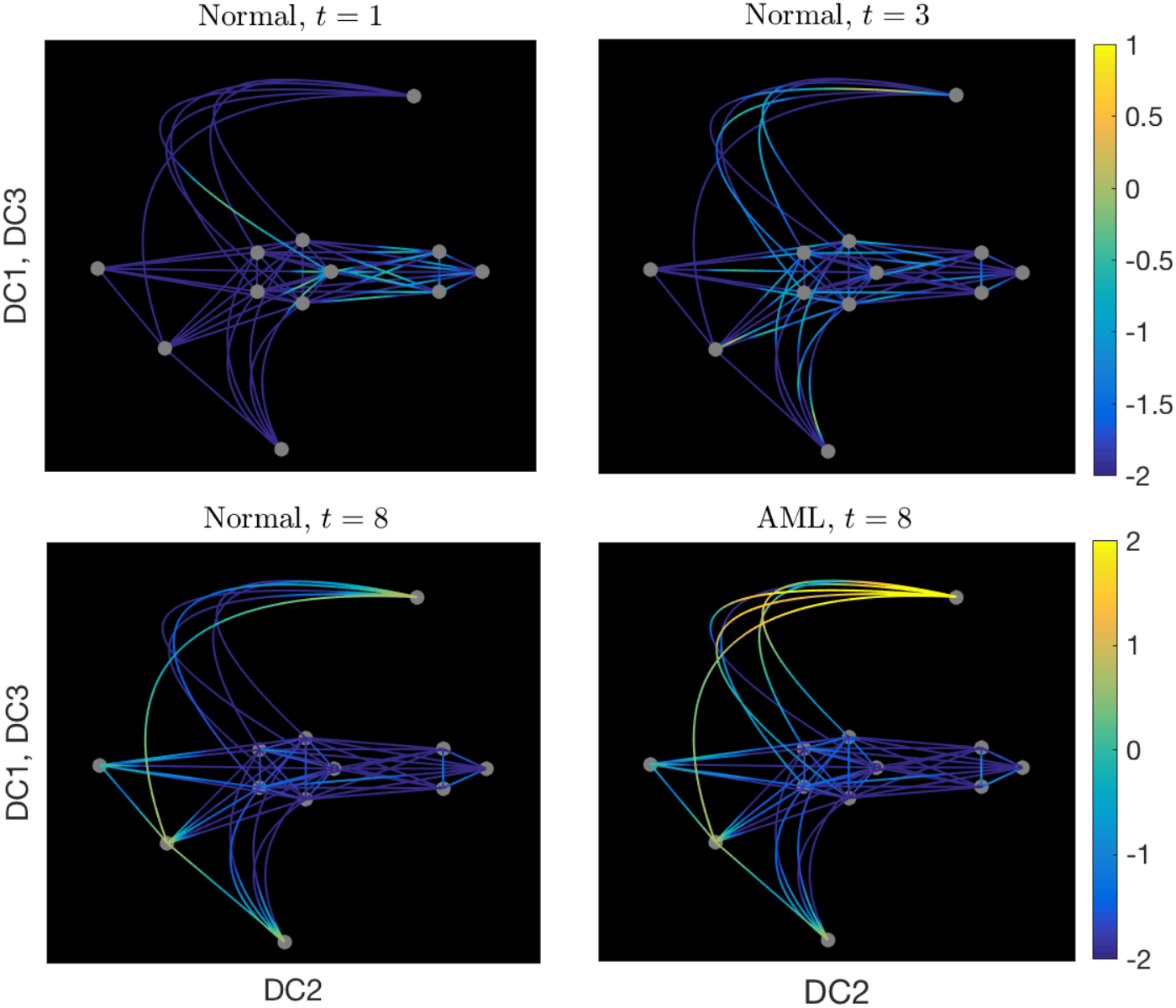
The cell distribution on the graph in a log_10_ scale, comparing the normal and AML conditions at *t* = 8. The AML condition shows increased density on the edges near the MEP state (node 11) at *t* = 8.

**Figure 8.**
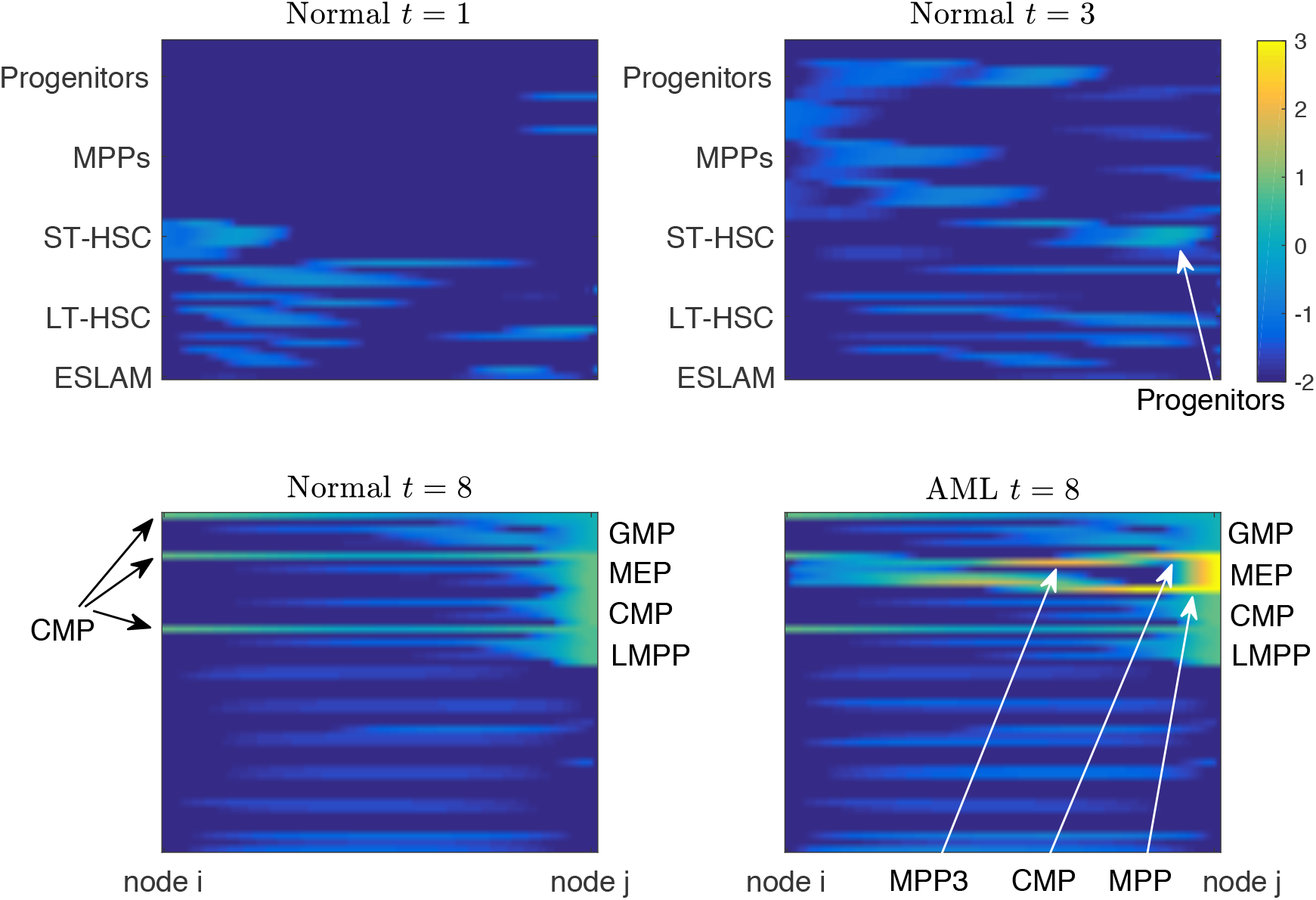
The continuum of cell states can be visualized as the density of cells along the 51 edges of the graph (rows) connecting node *i* (left) to node *j* (right) for all nodes *i, j*. Cell distribution (log_10_ scale) on the edge comparing the normal condition and AML. In addition to an accumulation of MEP cells, novel intermediate cell states emerge resulting from the differentation block and increased proliferation rate resulting from AML. These novel cell states are indicated with white arrows and generally fall between the CMP, MPP, and MEP canonical cell states.

We also simulated AML starting at different time points from *t* = 0 to *t* = 6. Since our initial condition assumes that the cells have not yet developed to MEP, the total number of cells is maximized when the AML occurs after a critical amount of cells have differentiated into an MEP state. Figure 9 shows the results of model simulations, where we observe that the number of cells are maximal at later times when AML is started at *t* = 3. From these simulations, we infer that the short and long term evolution of AML may depend on the state and composition of the hematopoietic landscape at the time of AML initiation.

**Figure 9.**
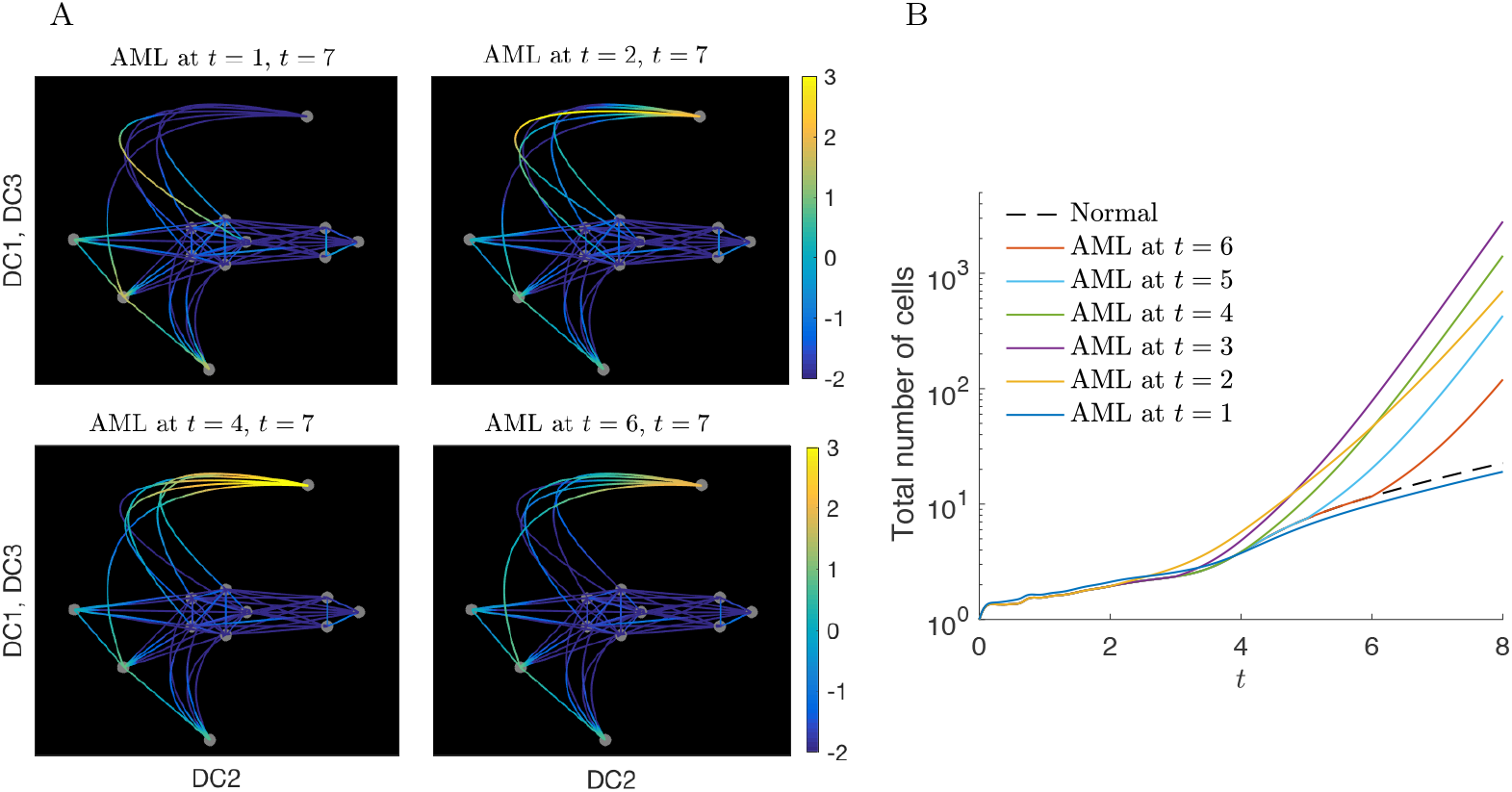
A) Cell distribution on the graph at *t* = 7 for AML occurring at different times, *t* = 1, 2, 4, and 6. MEP (11) blows up when AML occurs after *t* ≥ 2. The dominating intermediate cells are also distinct. B) Relative total number of cells when AML occurs at *t* = 1 to *t* = 6 compared to the normal case (dashed line) up to time *t* = 7. The total number of cells is maximized when AML occurs at *t* = 3. The bottom figure shows the cell distribution at *t* = 7 of different AML simulation starting at *t* = 1, 2, 4, and 6.

## 5. Discussion

We present a mathematical model of movement in an abstract space representing states of cellular differentiation. We represent trajectories in the differentiation space as a graph and model the directed and random movement on the graph with partial differential equations. We demonstrate our modeling approach on a simple graph, and then apply our model to hematopoiesis with publicly available scRNA-seq data. We calibrate the PDE model to pseudotime trajectories in the diffusion map space and use the model to predict the early stages of pathogenesis of acute myeloid leukemia.

Although methods exist to characterize differentiation trajectories, such as optimal transport (Schiebinger et al. 2017) and diffusion pseudotime methods (Haghverdi et al. 2016), an advantage of our approach is the ability to use a mathematical model to *predict* the outcomes of abnormal trajectories and to *perturb* the system mathematically with the model. We use this advantage of the mathematical model to simulate and explore AML pathogenesis based on immunophenotypic characterization of a mouse model for inv(16) AML. Our simulation results are consistent with the evolution of inv(16) driven AML, and predicts dynamics in canonical cell populations as well as cells in novel, intermediate states of differentiation. The intermediate cell states such as CMP-MEP seen in our simulation is consistent with previous observations that CBF*β*-SMMHC expressing phenotypic MEP cells confer CMP-like progenitor cell activity (Kuo et al. 2006). Given the phenotypic plasticity and aberrant differentiation occurring during leukemia evolution, it is particularly more informative to model cell dynamics in a continuum of differentiation space.

The novelty and power of this modeling approach is the ability to capture and predict dynamics of many interconnected cell types without the need to explicitly model cell types as discrete entities, which requires mathematical rules for their evolution and interactions. We now consider a continuum of cellular states, and model movement between these states in aggregate by representing many cell populations and states in a single variable. This approach increases biological resolution of the system by characterizing an infinite number of sub-states in a continuum representation and allows us to make predictions with one equation and very few model parameters, which can be directly calibrated to experimental data.

A limitation of this approach is that it does not include physical properties of the living biological system, such as the cellular micro-environment, which is known to play a critical role in the transformation of cell state and function. Furthermore, we recognize and acknowledge that cellular state transition dynamics as represented as a projection in a low dimensional space is an approximation of the dynamics in the original high dimensional space. Moreover, the dynamics observed and predicted in the lower dimensional space critically depend on the method of dimension reduction. This logic motivates our use of diffusion maps as the method to construct the differentiation space.

However, despite these limitations, we contend that this kind of analysis is a critical and valuable first step towards understanding the evolution of the higher dimensional system, and that low dimensional approximations have value, particularly when predictions in the lower dimensional space can be experimentally validated. We postulate that when dynamics in low dimensional representations are sufficiently characterized, they may eventually be used as a surrogate for high dimensional data, thus reverting the trend of “big data” back down to more informative “small data.”

We note that our modeling approach can be applied to any data set or manifold shape. As more normal and abnormal cellular state transitions are characterized at single cell resolution, we may apply similar computational and modeling methods to those systems. We emphasize our modeling approach is general and is not tailored or adapted to hematopoiesis in particular. Future applications of this approach may be useful to model the effects of therapies which target specific states of differentiation or the differentiation process itself.

## Acknowledgements

Research reported in this publication was supported by the National Cancer Institute of the National Institutes of Health under award number P30CA033572 and R01CA178387. The content is solely the responsibility of the authors and does not necessarily represent the official views of the National Institutes of Health.

## Disclosure statement

No potential conflicts of interest are disclosed by the authors.

## Supplementary figures

**Figure 1.**
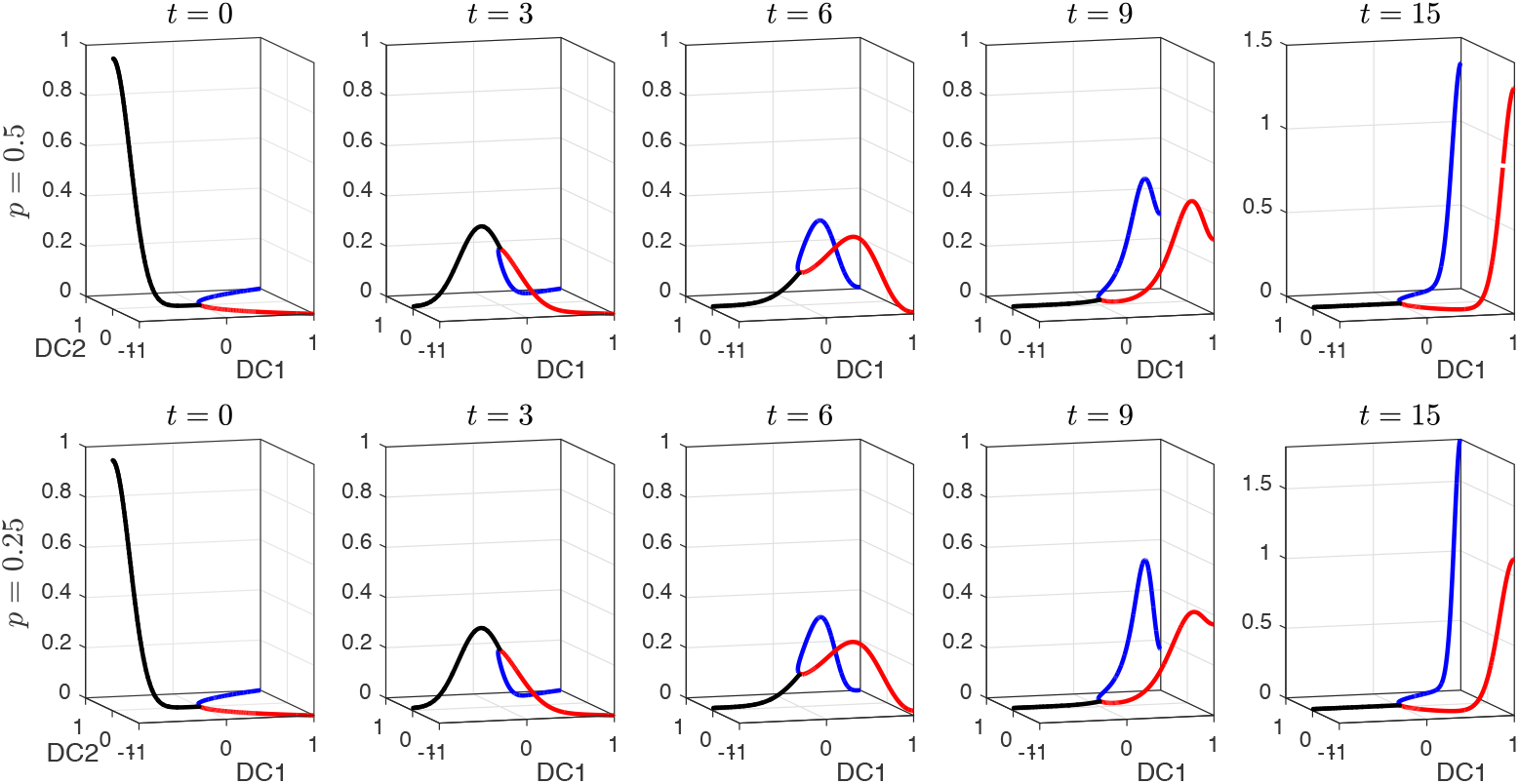
Solutions of the PDE model on the Y shaped graph from the initial condition centered at the left end *x* = −1 (black line in (a-b)) with diffusion *D* = 10^−2^ and drift *c* = −0.2 for the symmetric (top row) and asymmetric cases (bottom row).

**Figure 2.**
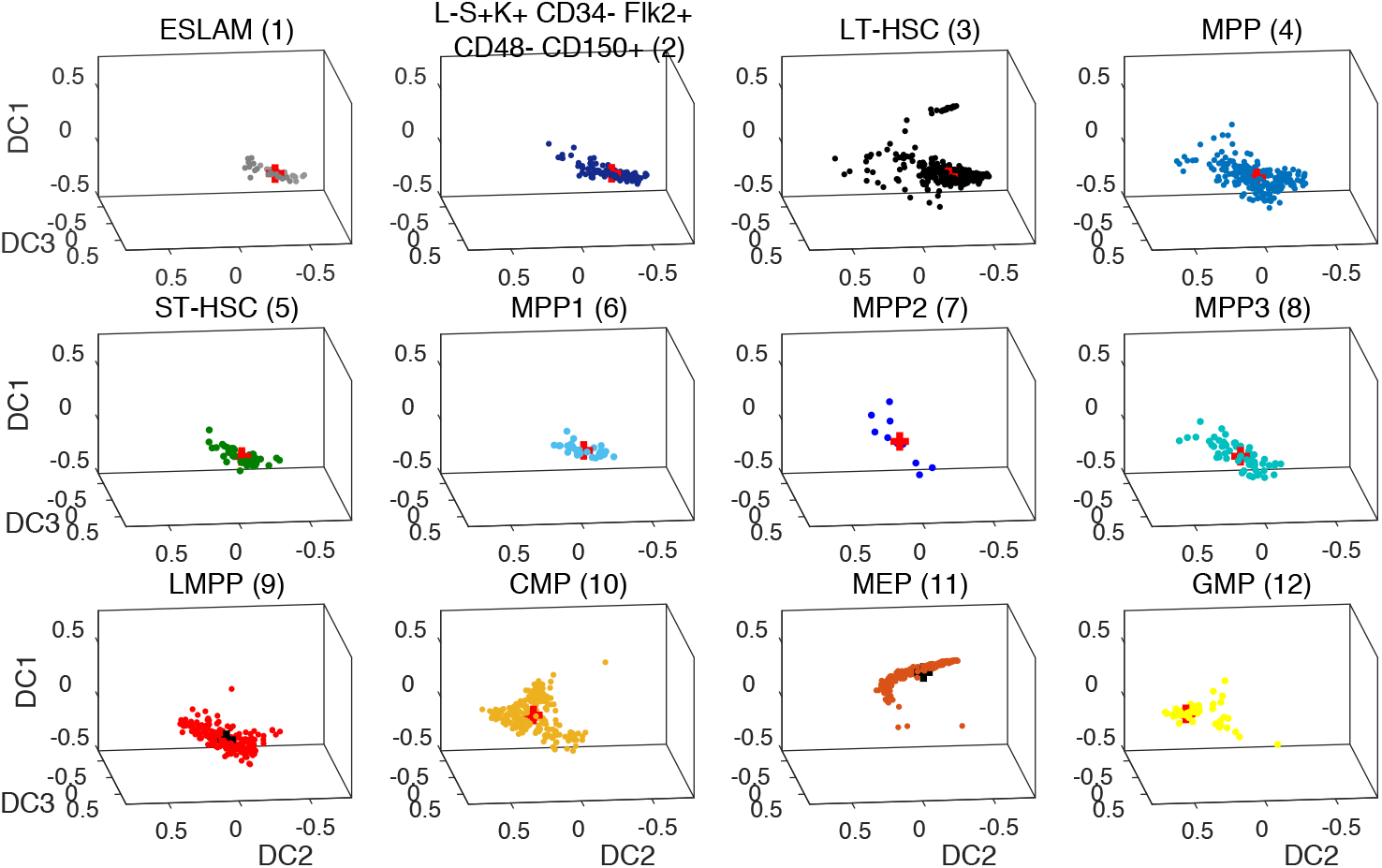
The cell data from (Nestorowa et al. 2016) is grouped into 12 cell nodes according to 12 commonly sorted HSPC phenotypes including LT-HSC, ST-HSC, and MEP. The center of mass of each cluster is marked as a red cross and used to establish nodes and edges on the graph which is then used as a computational domain for our simulations.

